# TMEM106B C-Terminal fragment aggregation exacerbates α−synuclein and TDP-43 proteinopathy phenotypes

**DOI:** 10.64898/2026.10.29.762961

**Authors:** Ruben Riordan, Aileen Saxton, Marina Han, Kailee Parrott, Brian Kraemer

## Abstract

Genetic variation in the human gene encoding the lysosomal transmembrane protein 106B (TMEM106B) drives risk of neurodegenerative dementia. A C-terminal fragment of TMEM106B (TMEM CT) accumulates in lesions across neurodegenerative disease, potentially contributing to neurotoxicity and hampering efficacy of therapeutic interventions. Previously, we developed the first transgenic organism model of TMEM CT neuronal aggregation in *Caenorhabditis elegans* (*C. elegans*). We demonstrated that TMEM CT neuronal aggregation drives severe neuronal dysfunction and neuronal loss. To investigate the potential synergistic interaction of TMEM CT aggregation with other types of aggregated neurotoxic proteinopathies occurring in dementia, we assessed neuronal degeneration and pathogenic protein accumulation in bigenic *C. elegans* expressing TMEM CT and other key proteinopathies of neurodegeneration including a-synuclein, tau, and TAR DNA binding protein 43 (TDP-43). Our results showed strong neurotoxic synergy of TMEM106B with TDP-43 or α-synuclein. However, we detected only mild synergy between TMEM106B and tau. This work provides evidence in support of TMEM CT aggregation as a significant participant in the neurodegenerative process attributed to other toxic protein aggregates. The work also highlights the importance of considering secondary proteinopathy contributions including TMEM CT aggregation when developing interventions against primary TDP-43 or α-synuclein proteinopathies.

**Summary Statement:** This manuscript assesses the potential synergistic impact of neuronal TMEM106B C-terminal aggregation on hallmark drivers of neurodegenerative diseases like TAR DNA binding protein 43, α-synuclein, or tau.

## Introduction

Transmembrane protein 106B (TMEM106B) is a type II lysosomal transmembrane protein governing locomotion and activity of lysosomes, primarily in neurons and oligodendrocytes (Feng, Lacrampe and Hu, 2021). Single nucleotide polymorphisms (SNPs) in TMEM106B have been identified as risk factors for various neurodegenerative diseases, most significantly in cases of frontotemporal lobar degeneration (FTLD) driven by TAR DNA binding protein 43 (TDP-43). However, as most of these TMEM106B SNPs are non-coding, the field struggled to identify the molecular mechanisms contributing to disease pathology (Nicholson *et al*., 2013; Brady, Zhou and Hu, 2014; Feng, Lacrampe and Hu, 2021; Hu *et al*., 2021; Perez-Canamas *et al*., 2021; Edwards *et al*., 2024; Marks *et al*., 2024). Recent analysis of post-mortem brain tissue from human cases of various neurodegenerative diseases, including Alzheimer’s Disease (AD), FTLD, progressive supranuclear palsy (PSP), and Lewy Body Dementia (LBD) showed the C-terminal fragment of TMEM106B (TMEM CT) is aggregating cytoplasmically in these diseases (Chang *et al*., 2022; Fan *et al*., 2022; Jiang *et al*., 2022; Hoq *et al*., 2023; Jiao, Yuan and Yu, 2023; Neumann *et al*., 2023; T Vicente *et al*., 2023; Bacioglu *et al*., 2024; Zhao *et al*., 2024).

Under normal conditions, TMEM CT, through glycosylation at sites N184 and N256, directs the protein to the lysosomal membrane. The TMEM CT sits on the luminal side of the membrane, where it is cleaved and lysosomally degraded (Feng, Lacrampe and Hu, 2021). It appears that with age, and in disease, TMEM CT is avoiding its lysosomal fate, resulting in cytoplasmic aggregation. Our previous work, developing the first and currently only live model of TMEM CT aggregation, demonstrated that TMEM CT aggregation alone is sufficient to drive severe neurotoxicity and reduced lifespan (Riordan *et al*., 2025). In human cases of dementia, TMEM CT appears to aggregate along with the hallmark proteins of neurodegenerative disease like tau, α-synuclein, and TDP-43 (Chang *et al*., 2022; Fan *et al*., 2022; Jiang *et al*., 2022; Hoq *et al*., 2023; Jiao, Yuan and Yu, 2023; Neumann *et al*., 2023; T Vicente *et al*., 2023; Bacioglu *et al*., 2024; Zhao *et al*., 2024). This raises the question as to whether or not TMEM CT aggregation is synergistically driving neurotoxicity with these pathogenic proteins in human disease.

A frequent occurrence in neurodegenerative disease is the accumulation of co-occurring protein pathologies, or multi-proteinopathies. AD is a disease characterized by co-pathology of tau and amyloid-β (Aβ), and is often complicated by the addition of Limbic Associated TDP-43 Encephalopathy (LATE), characterized by dysfunction and aggregation of TDP-43 (Braak *et al*., 2011; Nelson *et al*., 2019, 2024; Busche and Hyman, 2020; Kapasi *et al*., 2020). Further, neurodegenerative diseases that are characterized by a single pathological protein like FTLD or amyotrophic lateral sclerosis (ALS) with TDP-43 frequently coincide with other diseases driven by different pathogenic proteins like α-synuclein and tau (Körner *et al*., 2013; Trojsi *et al*., 2017; Chornenkyy, Fardo and Nelson, 2019; Latimer and Liachko, 2021; Moda *et al*., 2023; Tomé *et al*., 2023; Jellinger, 2026). In these diseases, little is known about how co-occurring protein pathologies interact to promote neurotoxicity as well as how or if therapeutically targeting one protein affects pathology of the other. For example, if a therapeutic approach targeting tau mediated pathology does not impact accumulation of pathogenic α-synuclein, this approach will have limited efficacy for patients experiencing co-pathology of AD and LBD. This gap in our knowledge is a major roadblock against the development of effective therapeutic interventions against a variety of neurodegenerative diseases, and is further complicated by the fact that TMEM CT is frequently co-aggregating in patients across a wide range of neurodegenerative diseases (Chang *et al*., 2022; Fan *et al*., 2022; Jiang *et al*., 2022; Hoq *et al*., 2023; Jiao, Yuan and Yu, 2023; Neumann *et al*., 2023; T Vicente *et al*., 2023; Bacioglu *et al*., 2024; Zhao *et al*., 2024).

The recent findings that TMEM CT aggregation on its own can drive neurotoxicity, and frequently co-aggregates with typical dementia related proteins across a wide range of neurodegenerative diseases suggests TMEM CT driven pathology may also pose a challenge to current attempts to develop therapeutic interventions against neurodegenerative disease (Riordan *et al*., 2025). As such, understanding the potential of TMEM CT aggregation to synergistically promote neurotoxicity when it coincides with other protein pathologies has become imperative. Specifically with TDP-43, multiple studies have already indicated that levels of TMEM CT aggregation may impact severity of TDP-43 driven pathology in human patients and impact seeding potential of TDP-43 aggregates isolated from human tissue (Mao *et al*., 2021; Zhong *et al*., 2025; Dopler *et al*., 2026).

To investigate the potential of TMEM CT aggregation to synergistically promote neurotoxicity when co-expressed with hallmark drivers of neurodegeneration, we conducted genetic analysis in bigenic *C. elegans* models pan-neuronally co-expressing low levels of TMEM CT aggregation in combination with TDP-43, α-synuclein, polyglutamine expansion repeats of 40 in the huntingtin protein (polyQ_40_), tau, or Aβ. With these bigenic models, we assessed changes to neuronal function, neurotoxicity, and pathogenic protein accumulation. We found that TMEM CT aggregation had noticeable synergy with both neuronal function and pathogenic protein accumulation when co-expressed with TDP-43, α-synuclein, and polyQ_40_, suggesting a shared mechanism of toxicity. TMEM CT aggregation showed a synergistic impact on neuronal dysfunction with pathogenic tau, but not on pathogenic protein accumulation, suggesting independent mechanisms of toxicity.

## Results

### Impact of TMEM CT aggregation on tau driven protein pathology and neuronal toxicity

Tau aggregation is a primary driver of neurodegeneration in AD and related dementia disorders including PSP (Braak *et al*., 2011; Cristiani *et al*., 2024). SNPs in TMEM106B are known to confer risk to both of these diseases, and aggregation of TMEM CT has been directly observed in the brains of patients with AD and PSP (Chang *et al*., 2022; Jiao, Yuan and Yu, 2023). However, it remains unknown whether TMEM CT aggregation can impact tau pathology to exacerbate neurotoxicity and disease progression. To test the impact of TMEM CT aggregation on tau driven pathology, we conducted a genetic cross between a low neuronally expressing TMEM CT *C. elegans* strain with a strain modestly overexpressing human tau in neurons. With this bi-genic model, we assayed neuronal function, neuronal loss, and pathogenic protein accumulation. On their own, the low expressing TMEM CT strains accumulate significantly less TMEM CT aggregates and show no observable neuronal dysfunction (**Supplemental Figure 1, Table 1**).

**Table 1.**
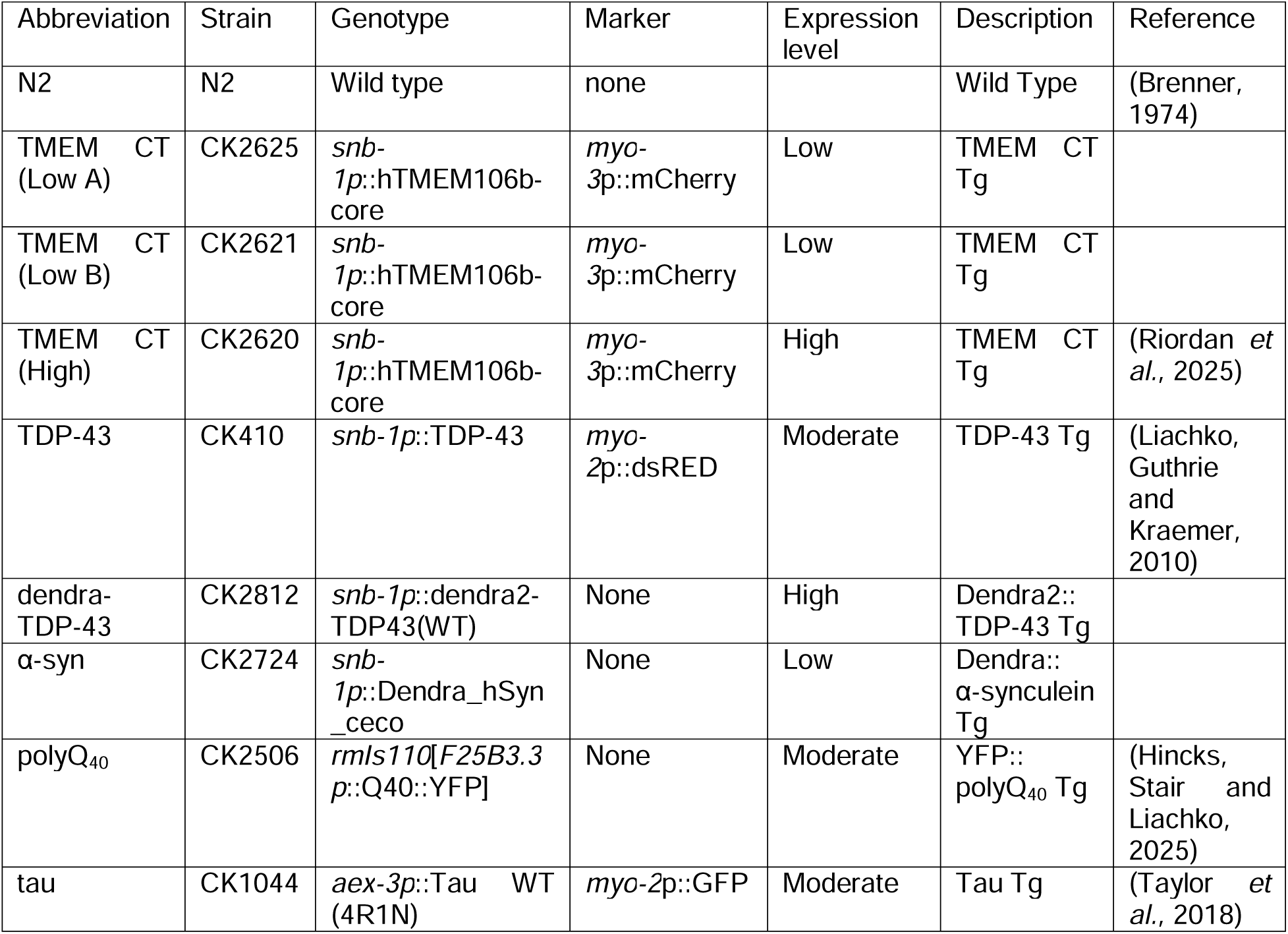
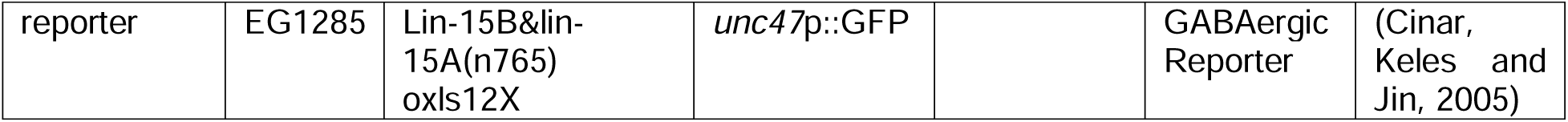
Strain List. List of *C. elegans* strains, genotypes, markers, descriptions, references, and abbreviations.

Western blot analysis of detergent insoluble protein fractions treated with formic acid (FA) shows that the presence of pathogenic tau did not impact levels of detergent insoluble TMEM CT. Similarly, analysis of detergent soluble samples shows that low aggregation of TMEM CT did not impact accumulation of either tau or phosphorylated tau (p-tau) (**Figure 1A**). Together, this indicates no direct interaction between accumulation of pathogenic tau and TMEM CT.

**Figure 1.**
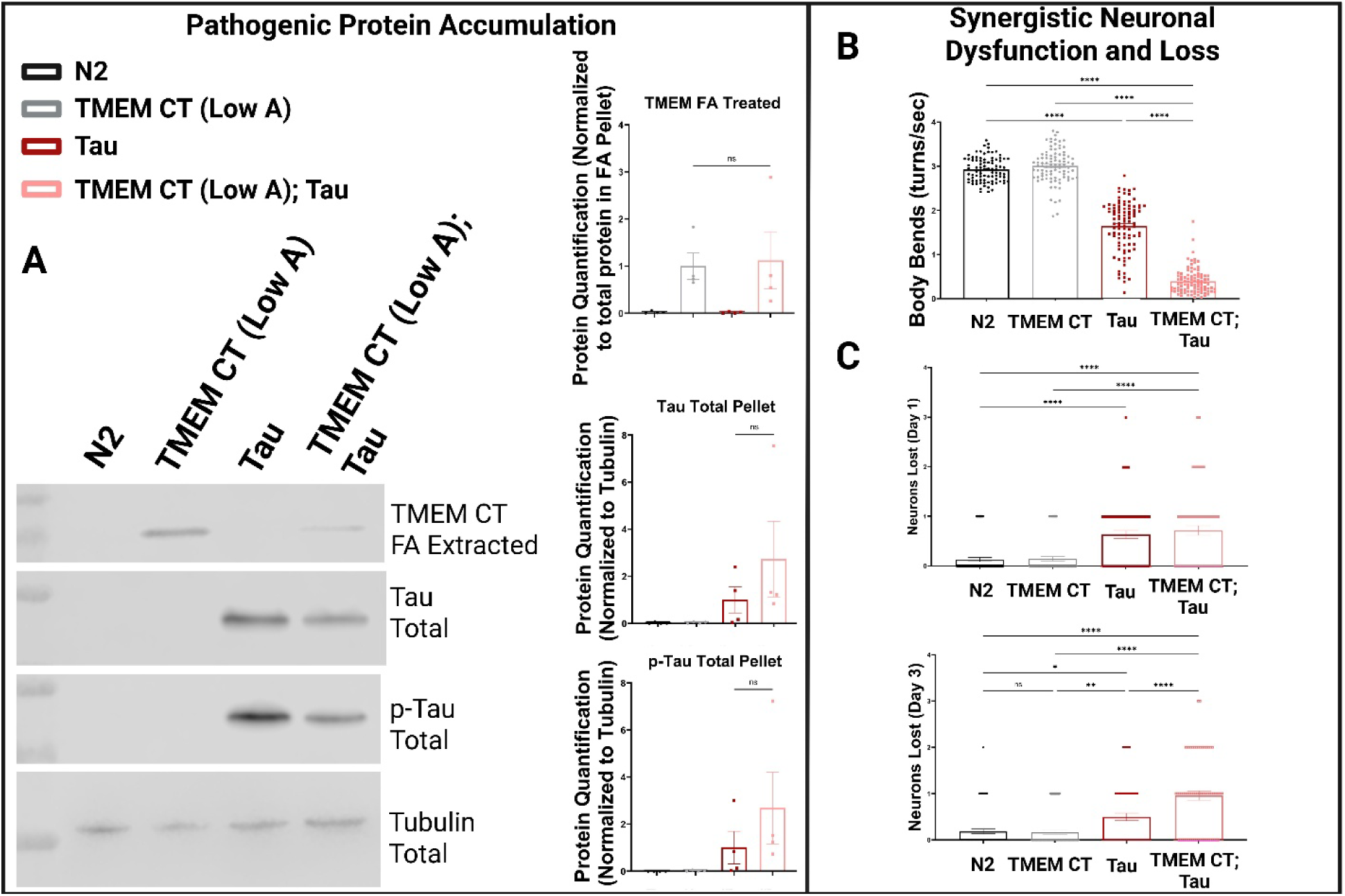
Impact of TMEM CT aggregation on Tau driven pathology. Comparison of non-transgenic *C elegans* (N2), single transgenic *C elegans* (TMEM CT (Low A) or Tau), and bigenic *C. elegans* (TMEM CT (Low A); Tau). **A)** Western blot analysis and quantification of *C. elegans* populations against TMEM CT (SYSY TMEM106B – 506 017) for detergent insoluble / FA treated samples and tau (Thermo Fisher SP70) or pTau (Hoofnagle Lab pT231) for detergent soluble protein samples. N = 4 for each strain, ** = P < 0.001. **B)** Liquid thrashing assay of motor neuron function at day 1 of adulthood, measuring body bends for each worm over a 60 s period. n > 60, N = 3 for each strain, **** = P < 0.00001. **C)** Counts of GABAergic neurons lost at day 1 or day 3 of adulthood using the *unc47*p:GFP reporter (EG1285). n > 60, N = 3 for each strain, **** = P < 0.00001.

To assess neuronal function in our bigenic model, we used a liquid based motility behavior assay (Kow *et al*., 2022). On day 1 of adulthood, we observed that our bigenic *C. elegans* expressing both pathogenic proteins had drastically impaired motor neuron function as compared to either single transgenic model expressing TMEM CT or Tau alone (**Figure 1B**). This indicates that TMEM CT and tau work synergistically to impair neuronal function. To assess changes to neuronal loss, we utilized the an *unc-47* promoter driven :GFP reporter *C. elegans* strain expressing GFP in all 19 GABAergic neurons along the nerve cord (Cinar, Keles and Jin, 2005). At day 3 of adulthood, but not day 1 of adulthood, we observed that animals expressing both TMEM CT and TDP-43 together lost significantly more GABAergic neurons as compared to the tau single transgenic model, indicating a synergistic impact on neuronal death between pathogenic TMEM CT and tau (**Figure 1 C**).

As TMEM106B SNPs are a risk factor for AD, TMEM CT aggregation has been observed in post-mortem brain tissue of AD patients, and aggregation of amyloid β 1-42 (Aβ_1-42_) is a pathological definer of AD in addition to tau pathology (Braak *et al*., 2011; Busche and Hyman, 2020), we were also interested in a potential interaction between TMEM CT aggregation and pathogenic Aβ_1-42_. To investigate this, we conducted a genetic cross between *C. elegans* neuronally expressing TMEM CT with *C. elegans* neuronally expressing Aβ_1-42_. However, we observed no interaction between TMEM CT and Aβ_1-42_ on neuronal function at day 1 or day 4 of adulthood (**Supplemental Figure 2**). It is important to note that the expression of Aβ_1-42_in this transgenic model is temperature sensitive. Regardless, we saw no impact on neuronal function when animals were incubated at 16, 20, or 25 degrees C.

### Impact of TMEM CT aggregation on polyQ_40_ protein pathology and neuronal dysfunction

The *HTT* gene, encoding the Huntington protein includes expansions of CAG repeats, translating into polyglutamine (polyQ) tracts. The normal Q tract length of the Huntington protein ranges from 6 – 35 repeats. Huntington protein with 36 or more polyQ repeats becomes progressively aggregate prone, and is the pathogenic driver of Huntington’s Disease (HD) (Waldvogel *et al*., 2015). While HD is known to also involve other proteinopathies, like TDP-43 pathology (Chánez-Cárdenas and Vázquez-Contreras, 2012; Corrochano *et al*., 2012; Bai *et al*., 2024; Nguyen *et al*., 2025), no previous analysis has been done to investigate the potential prevalence or impact of TMEM CT co-aggregation with pathogenic polyQ repeats. To address this gap in knowledge, we conducted genetic crosses to create bigenic models neuronally expressing both human TMEM CT and Huntington protein with either 40 (polyQ_40_) glutamine repeats. Independent of TMEM CT pathology, polyQ_40_ models typical HD pathology (Rüb *et al*., 2016; Pradhan *et al*., 2022). With this model we assessed pathogenic protein accumulation and neuronal function as compared to both single transgenic models.

Western blot analysis of detergent insoluble protein fractions treated with formic acid (FA) shows that worms expressing both TMEM CT and polyQ_40_ as opposed to just TMEM CT appear to experience increased levels of TMEM CT aggregation. While analysis of detergent soluble samples did not show an increased accumulation of polyQ_40_, aggregation of TMEM CT did give rise to polyQ_40_ accumulating in the detergent insoluble protein fraction (**Figure 2A**). Together, this indicates that TMEM CT aggregation synergistically increases the accumulation of pathogenic polyQ_40_ and vise-versa.

**Figure 2.**
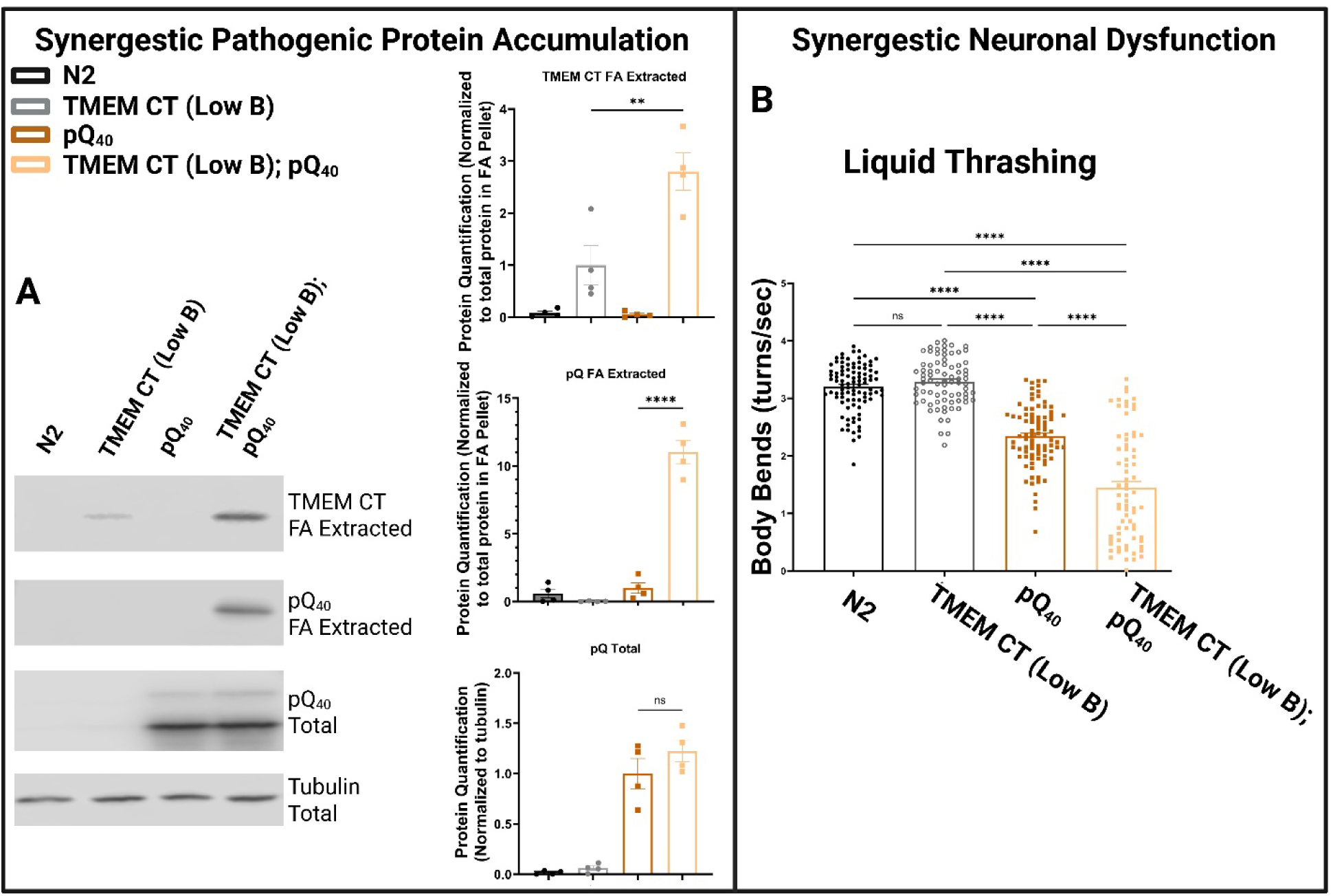
Impact of TMEM CT aggregation on polyQ_40_ driven pathology. Comparison of non-transgenic *C elegans* (N2), single transgenic *C elegans* (TMEM CT (Low A) or polyQ_40_), and bigenic *C. elegans* (TMEM CT (Low A); polyQ_40_. **A)** Western blot analysis and quantification of *C. elegans* populations against TMEM CT (SYSY TMEM106B – 506 017) and yfp for detergent insoluble / FA treated samples and yfp (Rockland anti-GFP) for detergent soluble protein samples. N = 4 for each strain, ** = P < 0.001 and **** = P < 0.00001. **B)** Liquid thrashing assay of motor neuron function at day 1 of adulthood, measuring body bends for each worm over a 60 s period. n > 60, N = 3 for each strain, **** = P < 0.00001.

Our polyQ expansion models express polyQ_40_ fused to a YFP tag, allowing for fluorescent recovery after photo-bleaching (FRAP) experiments to further test protein solubility of our polyQ expansions in live animals. FRAP analysis of *C. elegans* expressing both TMEM CT and polyQ_40_ showed drastically less recovery of fluorescent signal to the ablated area as compared to *C. elegans* expressing polyQ_40_ alone (**Supplemental Figure 3**). This indicates that polyQ_40_ is less mobile and therefore more aggregated in the presence of TMEM CT aggregation, supporting our western blot analysis.

To assess neuronal function in our bigenic model, we used a liquid thrashing assay. At day 1 of adulthood, we observed that our bigenic *C. elegans* expressing both pathogenic TMEM CT and polyQ_40_ had drastically impaired motor neuron function as compared to either single transgenic model expressing TMEM CT or polyQ_40_ alone (**Figure 2B**). This indicates that TMEM CT and polyQ_40_ work synergistically to impair neuronal function.

### Impact of TMEM CT aggregation on α-synuclein protein pathology and neuronal dysfunction

Aggregation of α-synuclein into Lewy bodies is the defining feature of DLB and Parkinson’s disease (Spillantini *et al*., 1998; Kim, Kågedal and Halliday, 2014; Kon, Tomiyama and Wakabayashi, 2020). While SNPs in TMEM106B are known to confer risk for both DLB and Parkinson’s Disease, and TMEM CT aggregation has been observed in human cases of both (Chang *et al*., 2022; Schweighauser *et al*., 2022), no previous analysis has been done to assess if there is any synergistic impact of TMEM CT and α-synuclein co-pathology on neurotoxicity. To address this gap in knowledge, we conducted a genetic cross to create a bigenic model neuronally expressing both human α-synuclein and low levels of TMEM CT. With this model we assessed pathogenic protein accumulation and neuronal function as compared to both single transgenic models.

Western blot analysis of detergent insoluble protein fractions extracted with formic acid (FA) shows that worms expressing both TMEM CT and α-synuclein as opposed to just TMEM CT experience drastically elevated levels of TMEM CT aggregation. While analysis of detergent soluble samples did not show an increased accumulation of α-synuclein, aggregation of TMEM CT did give rise to α-synuclein accumulating in the detergent insoluble protein fraction (**Figure 3A**). Together, this indicates that TMEM CT aggregation synergistically increases the accumulation of pathogenic α-synuclein and vice-versa.

**Figure 3.**
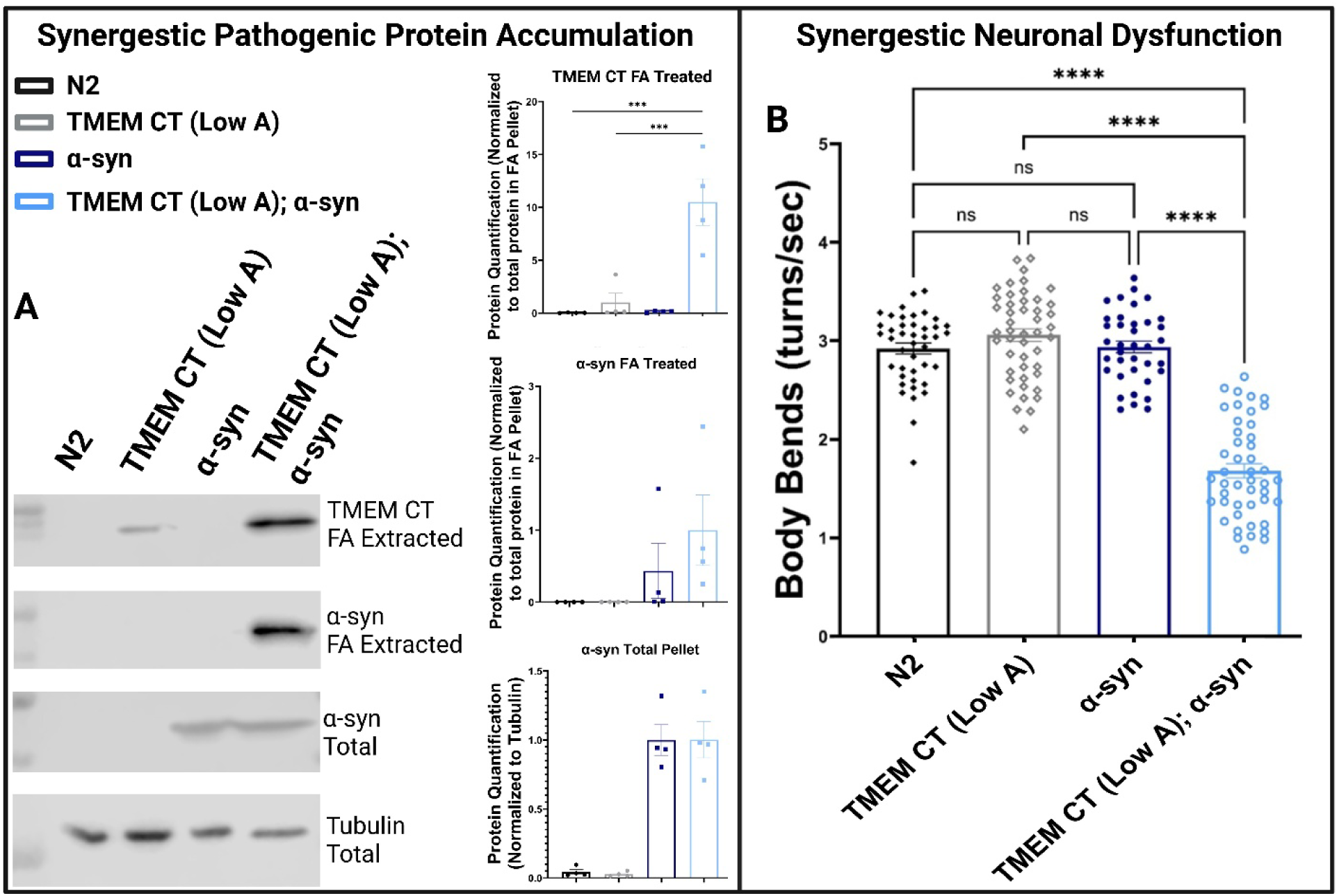
Impact of TMEM CT aggregation on α-synuclein driven pathology. Comparison of non-transgenic *C elegans* (N2), single transgenic *C elegans* (TMEM CT (Low A) or α-synuclein), and bigenic *C. elegans* (TMEM CT (Low A); α-synuclein). **A)** Western blot analysis and quantification of *C. elegans* populations against TMEM CT (SYSY TMEM106B – 506 017) and α-synuclein for detergent insoluble / Formic Acid extracted samples and α-synuclein (Thermo Fisher LB509) for detergent soluble protein samples. N = 4 for each strain, *** = P < 0.0001. **B)** Liquid thrashing assay of motor neuron function at day 1 of adulthood, measuring body bends for each worm over a 60 s period. n > 60, N = 3 for each strain, **** = P < 0.00001.

To assess neuronal function in our bigenic model, we used a liquid thrashing assay. At day 1 of adulthood, we observed that our bigenic *C. elegans* expressing both pathogenic proteins had drastically impaired motor neuron function as compared to either single transgenic model expressing TMEM CT or α-synuclein alone (**Figure 3B**). This indicates that TMEM CT and α-synuclein work synergistically to impair neuronal function.

### Impact of TMEM CT aggregation on TDP-43 protein pathology and neurotoxicity

TDP-43 cytoplasmic localization and aggregation drives neurodegeneration in ALS, FTLD-TDP, and LATE (Jo *et al*., 2020; Mackenzie and Neumann, 2020; Neumann *et al*., 2023; Young *et al*., 2023; Nelson *et al*., 2024). SNPs in TMEM106B are known to confer risk to each of these diseases, and abundant aggregation of TMEM CT has been directly observed in the brains of patients with ALS and FTLD-TDP (Mao *et al*., 2021; Jiang *et al*., 2022; Schweighauser *et al*., 2022; Marks *et al*., 2024; Dopler *et al*., 2026). Further, a recent study by Zhong et al. showed that levels of TMEM CT aggregates extracted along with TDP-43 aggregates from human cases of FTLD explained variability in TDP-43 seeding potential in human cell culture models (Zhong *et al*., 2025).

To test the impact of TMEM CT aggregation on TDP-43 driven pathology, we conducted a genetic cross between a low neuronally expressing TMEM CT *C. elegans* strain with a strain modestly overexpressing human TDP-43 in neurons. With this bi-genic model, we assayed neuronal function, neuronal loss, and pathogenic protein accumulation.

Western blot analysis of detergent insoluble protein fractions solubilized with formic acid (FA) show that worms expressing both TMEM CT and TDP-43 as opposed to just TMEM CT experience drastically elevated levels of TMEM CT aggregation. Similarly, analysis of non FA treated samples show that worms expressing both TMEM CT and TDP-43 exhibit increases accumulation of both total TDP-43 and phosphorylated TDP-43 (pTDP-43) when compared to TDP-43 Tg alone (**Figure 4A**). Together, this indicates that TMEM CT aggregation synergistically increases the accumulation of pathogenic TDP-43 species and vice versa.

**Figure 4.**
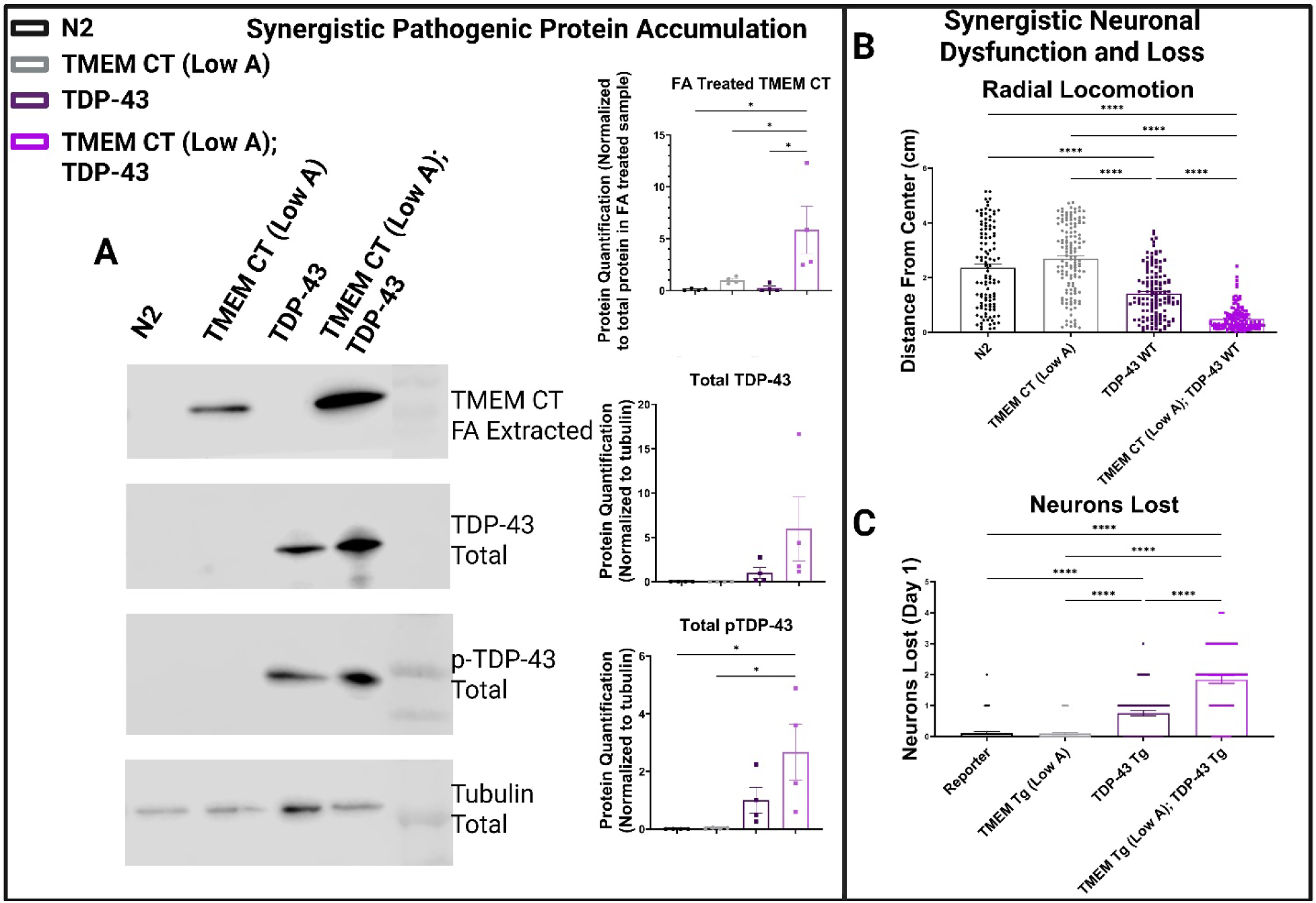
Impact of TMEM CT aggregation on TDP-43 driven pathology. Comparison of non-transgenic *C elegans* (N2), single transgenic *C elegans* (TMEM CT (Low A) or TDP-43), and bigenic *C. elegans* (TMEM CT (Low A); TDP-43). **A)** Western blot analysis and quantification of *C. elegans* populations against TMEM CT (SYSY TMEM106B – 506 017) for detergent insoluble / FA treated samples and TDP-43 (AB57105) or pTDP-43 (WSU Monoclonal Antibody Center 3C) for detergent soluble protein samples. N = 4 for each strain, * = P < 0.05. **B)** Radial locomotion assay of motor neuron function at day 1 of adulthood, measuring distance traveled for each worm over a 30 minute period. n > 60, N = 3 for each strain, **** = P < 0.00001. **C)** Counts of GABAergic neurons lost at day 1 of adulthood using the *unc47*p:GFP reporter (EG1285). n > 60, N = 3 for each strain, **** = P < 0.00001.

To assess neuronal dysfunction in our bigenic model, we used a radial locomotion assay. At day 1 of adulthood, we observed that our bigenic *C. elegans* expressing both pathogenic proteins had drastically impaired locomotion as compared to either single transgenic model expressing TMEM CT or TDP-43 alone (**Figure 4B**). This indicates that TMEM CT and TDP-43 work synergistically to impair neuronal function. To assess changes to neuronal loss, we utilized the GABAergic reporter described above. At day 1 of adulthood, we observed that animals expressing both TMEM CT and TDP-43 together exhibited significantly more GABAergic neurons as compared to either single transgenic model, indicating a synergistic impact increasing neuronal loss when both pathogenic proteins are present (**Figure 4C**).

## Discussion

Co-occurring distinct pathogenic proteinopathies in neurodegenerative disease appear to be more common than isolated single proteinopathy disorders.The predominance of multi-proteinopathy likely confounds attempts to developtruly effective therapeutic interventions against devastating diseases including AD, FTLD, PD, and ALS (Braak *et al*., 2011; Chánez-Cárdenas and Vázquez-Contreras, 2012; Trojsi *et al*., 2017; Wang *et al*., 2018; Chornenkyy, Fardo and Nelson, 2019; Nelson *et al*., 2019, 2024; Busche and Hyman, 2020; Kapasi *et al*., 2020; Huang *et al*., 2021; Latimer and Liachko, 2021; Latimer *et al*., 2022; Moda *et al*., 2023; Tomé *et al*., 2023; Bai *et al*., 2024; Jadhav *et al*., 2024; Zhao *et al*., 2024; Franzmeier *et al*., 2025; Gawor *et al*., 2025; Nguyen *et al*., 2025). In fact, co-pathology of pathogenic Aβ_1-42_ and tau defines AD. (Braak *et al*., 2011; Busche and Hyman, 2020). AD commonly co-occurs with LATE, adding TDP-43 pathology to the neurodegenerative process (Kapasi *et al*., 2020; Nelson *et al*., 2024). We now know that TMEM CT cytoplasmic aggregation occurs as a contributing neuropathological feature that manifests frequently across a wide range of neurodegenerative diseases (Chang *et al*., 2022; Fan *et al*., 2022; Jiang *et al*., 2022; Hoq *et al*., 2023; Jiao, Yuan and Yu, 2023; Neumann *et al*., 2023; T Vicente *et al*., 2023; Bacioglu *et al*., 2024; Zhao *et al*., 2024). Taken together these observations highlight the need to understand how TMEM CT aggregation interacts with common pathogenic proteins of neurodegeneration to promote neurotoxicity.

In this study, we make use of our previously described model of TMEM106B proteinopathy expressing TMEM CT in all neurons ((Riordan *et al*., 2025) and **Supplemental Figure 1**). With these low expressing TMEM CT strains, we conducted genetic analysis with transgenic *C. elegans* pan-neuronally expressing pathogenic human tau, Aβ_1-42_, polyQ_40_, α-synuclein, or TDP-43 to generate bigenic animals that model co-pathology of TMEM CT with hallmark drivers of neurodegenerative disease (Table 2).

**Table 2.** Impact of low neuronal TMEM CT aggregation on neurodegeneration driven by common drivers of dementia. Summary table indicating the strength (strong, moderate or no impact) of synergy between TMEM CT aggregation and common drivers of neurodegenerative disease on neuronal function, neuronal death, and pathogenic protein accumulation. * NA = not assessed.

|  | Neuronal Function | Neuronal Death | Protein Accumulation |
| --- | --- | --- | --- |
| **Tau** | Strong | Moderate | No Impact |
| **Aβ<sub>1-42</sub>** | No Impact | NA | NA |
| **polyQ<sub>40</sub>** | Strong | NA | Moderate |
| **α-syn** | Strong | NA | Strong |
| **TDP-43** | Strong | Strong | Strong |

We modeled the interaction between TMEM CT and tau to investigate the neurotoxic impact of TMEM CT secondary proteinopathy in tauopathy driven dementias, including AD and PSP (Braak *et al*., 2011; Cristiani *et al*., 2024). TMEM106B SNPs are a known risk factor for AD, and aggregation of TMEM CT occurs in the brains of patients with AD and PSP (Chang *et al*., 2022; Jiao, Yuan and Yu, 2023). However, prior to this study, no analysis has been conducted to evaluate the potential interactions between pathogenic tau and TMEM CT. With our bigenic model neuronally expressing both human tau and TMEM CT, we saw no impact of co-expression on accumulation of pathogenic tau or TMEM CT. Despite the lack of interaction on protein accumulation, assessment of neuronal function did show a synergistic impairment of motor neuron function at day 1 of adulthood. Counts of GABAergic neurons similarly showed a synergistic increase in neuronal death, though unlike with TDP-43, this synergy was not observed until day 3 of adulthood. As TMEM CT and tau neuronal co-expression results in increased neurotoxicity, but neither pathogenic protein impacts accumulation of the other, it appears that the impact on neuronal function was additive and their mechanisms of toxicity are independent. This supports our previous findings that strong genetic suppressors of tau pathology only have mild impacts on TMEM CT driven pathology (Riordan *et al*., 2025).

We then modeled the potential interaction between TMEM CT and polyQ expansion repeats in the Huntington protein, which is the main driver of HD (Waldvogel *et al*., 2015). It is known that HD frequently co-occurs with other neurodegenerative protein pathologies like TDP-43 and α-synuclein proteinopathy (Chánez-Cárdenas and Vázquez-Contreras, 2012; Corrochano *et al*., 2012; Bai *et al*., 2024; Nguyen *et al*., 2025). However, no analysis has been done to assess the potential interaction between TMEM CT and polyQ expansion repeats. With our bigenic model neuronally expressing both human TMEM CT and polyQ_40_, modeling less severe HD pathology, we showed that the presence of polyQ_40_ resulted in drastically elevated levels of detergent insoluble TMEM CT, and the aggregation of TMEM CT gave rise to detergent insoluble polyQ_40_ Further, FRAP analysis displayed that the presence of TMEM CT drastically reduced motility of polyQ_40_ in live neurons, indicating more aggregation of polyQ_40_. Assessing neuronal function, we showed that TMEM CT and polyQ_40_ synergistically impaired motor neuron function in our model. Together, this indicates a strong synergistic impact of TMEM CT aggregation and polyQ expansion repeats on pathogenic protein accumulation and neurotoxicity.

We next modeled the potential interaction between TMEM CT and α-synuclein, the characterizing pathogenic protein of PD and DLB (Spillantini *et al*., 1998; Kim, Kågedal and Halliday, 2014; Kon, Tomiyama and Wakabayashi, 2020). Similar to with FTLD and ALS, TMEM106B SNPs are a known risk factor for PD, and aggregation of TMEM CT occurs in the brains of patients with PD and DLB (Chang *et al*., 2022; Schweighauser *et al*., 2022). Emerging evidence suggests that TMEM CT burden is higher in cases of PD as compared to control cases (Zhao *et al*., 2024; Meng *et al*., 2026). However, nothing is currently known about how or if TMEM CT and α-synuclein pathology work together to promote neurotoxicity. With our bigenic *C. elegans* model neuronally expressing both human TMEM CT and α-synuclein, we observed that pathogenic α-synuclein increases the accumulation of detergent insoluble TMEM CT and the aggregation of TMEM CT gave rise to the accumulation of detergent insoluble α-synuclein. Assaying for motor neuron function, we saw that modest TMEM CT aggregation synergistically impaired neuronal function with α-synuclein at day 1 of adulthood, despite neither single-transgenic model exhibiting neuronal dysfunction independently. Together, this indicates a strong synergistic impact of TMEM CT aggregation on α-synuclein pathogenic protein accumulation and neurotoxicity. Given the ability of TMEM CT to increase burden of α-synuclein accumulation and vice-versa, these pathogenic proteins may share some mechanisms of neurotoxicity.

Finally, we modeled the potential interaction between TMEM CT and TDP-43, the characterizing pathogenic protein of FTLD-TDP, ALS, and LATE (Jo *et al*., 2020; Mackenzie and Neumann, 2020; Neumann *et al*., 2023; Young *et al*., 2023; Nelson *et al*., 2024). It is already known that SNP’s in TMEM106B confer significant risk for TDP-43 driven diseases like FTLD-TDP and ALS, and aggregation of TMEM CT has been observed in post-mortem human tissue from these diseases (Mao *et al*., 2021; Jiang *et al*., 2022; Schweighauser *et al*., 2022; Marks *et al*., 2024; Dopler *et al*., 2026). Emerging evidence already suggests that TMEM CT aggregation is correlated to TDP-43 nuclear depletion in human tissue and can burden of TMEM CT aggregation can impact seeding potential of TDP-43 (Mao *et al*., 2021; Zhong *et al*., 2025). With our bigenic *C. elegans* model neuronally expressing both human TMEM CT and TDP-43, we observed that pathogenic TDP-43 increases the accumulation of detergent insoluble TMEM CT and the aggregation of TMEM CT increases the accumulation of detergent soluble TDP-43 and pTDP-43. Modest aggregation of TMEM CT did not, however, impair the nuclear localization of TDP-43. Assaying for motor neuron function, we saw that modest TMEM CT aggregation (which independently did not impact neuronal function) synergistically elevated TDP-43 driven neuronal dysfunction at day 1 of adulthood. Further, we showed that TMEM CT aggregation significantly increased loss of GABAergic neurons at day 1 of adulthood. Together, this indicates a strong synergistic impact of TMEM CT aggregation on TDP-43 pathogenic protein accumulation and neurotoxicity. Given the ability of TMEM CT to increase burden of TDP-43 accumulation and vice-versa, these pathogenic proteins may share some mechanisms of neurotoxicity.

In summary, low levels of TMEM CT aggregation had strong synergistic impacts on neuronal toxicity and pathogenic protein accumulation or aggregation when neuronally co-expressed with TDP-43, α-synuclein, and polyQ expansion repeats. It is likely that these pathologies share mechanisms of toxicity with TMEM CT aggregation. Therefore, investigating the neurotoxic mechanisms of TMEM106B pathology will likely reveal potential therapeutic targets against neurodegenerative diseases driven by these pathogenic proteins. To a lesser extent, low TMEM CT aggregation also had an additive impact on tau driven neurotoxicity, but through an independent mechanism of toxicity. For tau driven neurodegenerative diseases that frequently involve co-pathology of TMEM CT aggregation, like AD and PSP, it is unlikely that targeting tau pathology will ameliorate TMEM CT aggregation. Given our previous findings that high levels of TMEM CT aggregation alone is sufficient to drive neuronal death, this will be an important point of consideration when developing therapeutic interventions against diseases like AD and PSP.

The results in this study confirm our previous findings that neuronal TMEM CT aggregation has significant potential to contribute to neurodegeneration in human age and disease. Further, even low levels of TMEM CT aggregation, which have no impact on neuronal function alone, can drive increased neurotoxicity when co-occurring with typical protein pathologies of neurodegenerative disease. This validates the hypothesis that TMEM106B aggregation is an important secondary proteinopathy that frequently occurs in dementia and exacerbates neurodegeneration. It is imperative that the field further assess TMEM106B as a driver of neurodegenerative disease and investigate how TMEM CT aggregation interacts with pathogenic proteins like TDP-43 and α-synuclein in human disease. In this study, we have developed the only live organism models that captures TMEM CT aggregation as a secondary proteinopathy which will be used to address this gap.

## Methods

### *C. elegans* strains and transgenics

*C. elegans* transgenic strains and information are listed in Supplemental Table 1.

### Plasmids

To generate transgenes expressing the C-terminal fragment of human TMEM106B driven by a pan-neuronal snb-1 promoter, a *C. elegans* codon optimized DNA sequence encoding the C-terminal fibrillar core fragment of wild type TMEM106B (amino acids 120-254 of NP_001127704) was purchased Integrated DNA Technologies (IDT). The parental Psnb-1 vector was previously constructed (Liachko, Guthrie and Kraemer, 2010) by inserting the snb-1 promoter sequences into the HindIII and BamHI sites of MCS I of plasmid pPD49.26 (a generous gift from Dr. A. Fire, Stanford University, Palo Alto, CA).

### Transgenics and strains

N2 (Bristol) was used as wild-type *C*. *elegans* and maintained as previously described (Brenner, 1974). The marker transgene *myo-3p*::mCherry was used at a concentration of 20 ng/μl as a co-injection marker as previously described (Liachko, Guthrie and Kraemer, 2010). The *snb-1p*::hTMEM106B-core transgene described above was microinjected into N2 at a concentration of 150 ng/μl and 75 ng/ μl respectively as previously described (Kraemer *et al*., 2003) to produce animals expressing TMEM106B and mCherry transgenes as extrachromosomal arrays. Extrachromosomal transgenes were integrated by exposing animals to UV radiation in a Stratalinker for 0.8 minutes. Progeny of irradiated animals were screened for 100% transmission of the transgene and integrated strains were backcrossed to N2 at least three times.

Strain EG1285 has an integrated P*unc-47*::GFP transgene expressing GFP in GABAergic neurons that clearly marks the cell bodies and axons of ventral cord VD and DD type inhibitory motor neurons (Cinar, Keles and Jin, 2005). All TMEM106B transgenic strains described here contain integrated transgenes carrying human TMEM106B-core encoding cDNAs and a Pmyo-3::mCherry coinjection marker. CK2625, CK2621 and CK2620 carry *snb-1p*::TMEM106B-core (TMEM CT (Low A), TMEM CT (Low B), and TMEM CT (High). Double mutants with TMEM106B expression were generated by crossing CK2625 or CK2621 with other strains carrying other transgenes or mutations of interest.

All strains were maintained at 20 °C on NGM plates seeded with OP50 *Escherichia coli* (Brenner, 1974). Genotypes were confirmed by PCR and sequencing for all strains with non-obvious phenotypes.

### Live *C*. *elegans* imaging

Live worms were mounted on a 4% agarose pad and immobilized with ∼50 mM sodium azide [Sigma] and covered with a glass slip. Representative images were acquired using a Nikon A1R confocal microscope with a 10x air objective or 40x or 100x oil immersion lens (Nikon USA, Melville, NY). Z-plane stacked images were flattened into a maximum intensity projection using ImageJ software.

### Immunohistochemistry

Day 1 adult worms were fixed in 1% formaldehyde solution and permeabilized by freeze cracking as described previously (Crittenden and Kimble, 1999). Fixed and permeabilized animals were counterstained with 300 nM 4,6-diamidino-2-phenylindole (DAPI) nuclear stain in PBS. Fixed whole animals were stained with anti-TMEM CTF monoclonal antibody (SYSY TMEM106B – 506 017) (Held *et al*., 2025; Replogle *et al*., 2026) at a dilution of 1:1,000. Alexa 488-conjugated anti-rabbit antibody (Invitrogen) and ALEXA 647-conjugated anti-mouse antibody was used as the secondary antibody at a dilution of 1:1000. 100 times magnification images were acquired by a Nikon A1R confocal microscope with a 100x oil immersion lens (Nikon USA, Melville, NY) or Andor Dragonfly 200 100x oil immersion lens (Andor UK, Bellfast) and processed in FIJI:imageJ.

### Behavioral analysis

NGM plates of day 1 adult *C*. *elegans* were flooded with 1 mL of M9 buffer (22 mM KH_2_PO_4_ monobasic, 42.3 mM Na_2_HPO_4_, 85.6 mM NaCl, 1 mM MgSO_4_), and swimming worms were pipetted onto a 35 mm unseeded NGM plate. Approximately 30 s following the addition of M9 buffer, worms were recorded swimming for 1 min at 14 frames per second. These videos were captured and analyzed with WormLab 2021 (MBF Bioscience). The frequency of body bends, or turns, as defined as a change in body angle of a least 20° from a straight line measured by the quarter points and midpoint of the worms, was quantified as a readout of locomotion. Worms tracked for less than 30 s were omitted from this analysis. At least three independent samples totaling at least 60 worms per strain were counted for every comparison (Kow *et al*., 2022).

### Neurodegeneration assays

Worms at larval stage 4 (L4) were selected and moved onto new NGM plates with OP-50. 24 hours later, live worms were placed on a 2% agarose pad containing 0.01% sodium azide to immobilize the worms. Worms were imaged under fluorescence microscopy and scored for number of GABAergic neurons. At least three independent samples totaling at least 60 worms per strain were counted for every comparison. Data were analyzed using GraphPad Prism software (Saxton and Kraemer, 2021; Jin *et al*., 2022).

### Production of pS409/410 TDP-43 monoclonal antibody

A synthetic peptide corresponding to the c-terminus of TDP-43 with phosphoserines substituents at positions 409/410 were used to immunize mice to produce monoclonal antibodies in the Washington State University Monoclonal Antibody Center using standard operating procedures (Elnaggar *et al*., 2017). Monoclonal antibodies were produced at WSU as a fee for service project (IACUC protocol #6317) using well established standard operating protocols for antibody production (Elnaggar *et al*., 2017). Briefly, peptides were mixed in Sigma Adjuvant System at 50% (w/v) as an emulsion prior to intraperitoneal immunization as recommended by the manufacturer (Sigma Chemical, catalog S6322). Mice were immunized with 200 µg of peptide and received an identical boost immunization 3 weeks after initial immunization. Spleens were harvested 3 days after boosting for hybridoma production. Clonal hybridoma lines were screened based on binding activity for phosphorylated c-terminal peptide but the otherwise identical, but non-phosphorylated peptide and the pS409/410 clone 3C hybridoma monoclonal line was isolated and cloned.

### Protein extraction

*C*. *elegans* were grown from hypochlorite-purified eggs at 20 °C for 3 days on 5XPEP plates until young adults. Worms were washed off plates with M9 buffer, collected by centrifugation, and grown further on egg plates as previously described (Portman, 2006). Well-fed worms were collected from egg plates, harvested and isolated via sucrose floatation, washed into M9 buffer, and collected in 250 µL aliquots of packed worms prior to snap-freezing in liquid nitrogen to be stored at −70 °C. These worm samples were thawed, sonicated (11 x 15 s at 70% power and put on ice in between sonications) in TM-LS Buffer (10 mM Tris, 800 mM NaCl, 1 mM EDTA, 10% Sucrose, pH 7.5). Sarkosyl insoluble material was separated from soluble protein through ultra-centrifugation (2 spins at 140,000 x g for 30 mins each). The detergent insoluble material is not readily detectable without rigorous solubilization so a modified Aβ extraction protocol (Casali and Landreth, 2016) was utilized to expose TMEM epitopes. Briefly, aggregated protein was resolubilized by resuspension and sonication in 90% formic acid (FA) to expose TMEM fibrillar epitopes. FA samples were dried in a speedvac to evaporate off formic acid. Dried protein aggregate pellets were solubilized in 100 μL 5xSDS (46 mM Tris, 5 mM EDTA, 200 mM dithiothreitol, 50% sucrose, 5% sodium dodecyl sulfate, 0.05% bromophenol blue) protein sample buffer immediately after formic acid treatment. 200 μL aliquots were taken after initial solubilization in TM-LS buffer for soluble protein fractions and pelleted. Soluble protein was diluted with 60 μL 1× SDS protein sample buffer. Samples were sonicated at 70% for 15 s three times returning to ice between each replicate, boiled for 10 min at 95 °C, centrifuged at 13,000 × g for 2 min, and stored at −80 °C.

Samples were loaded (6 μL) onto 4-15% precast criterion sodium dodecyl sulfate polyacrylamide gel electrophoresis gradient gels and transferred to PVDF per manufacturer’s protocol (Bio-Rad). The protein sizing ladder used was Precision Plus Protein Standards (Bio-Rad). The primary antibodies used were rat monoclonal anti-TMEM106B C-terminus antibody (SYSY TMEM106B – 506 017) (Held *et al*., 2025; Replogle *et al*., 2026) at 1:5,000, mouse anti-tubulin antibody E7 (Developmental Studies Hybridoma Bank) at 1:5,000, rabbit monoclonal anti-tau antibody SP70 (Thermo Fisher) (Sanchez *et al*., 2021) at 1:5,000, rabbit monoclonal anti-ptau antibody pT231 (Hoofnagle lab) (Trivedi *et al*., 2025) at 1:5,000, mouse monoclonal anti-tdp-43 antibody AB57105 (Abcam) (Liachko *et al*., 2016) at 1:5,000, monoclonal mouse anti-TDP-43 phosphorylated at positions 409/410 antibody 3C (developed in collaboration with Washington State University Monoclonal Antibody Center (described above) at 1:10,000, mouse monoclonal anti-α-synuclein antibody LB509 (Invitrogen) (Israeli and Sharon, 2009) at 1:500, and goat polyclonal anti-GFP antibody (Rockland) at 1:2,000 (Birnbaum *et al*., 2023). The secondary antibodies used were anti-mouse HRP, anti-rabbit HRP, or anti-rat HRP (Jackson Immuno Research) at 1:5,000. ECL substrate was used to visualize the membrane (Bio-Rad). Chemiluminescence signals were detected with the LiCor Imager and quantified with Fiji (Schindelin *et al*., 2012). Insoluble protein samples were also loaded (6 μL) onto 4-15% precast criterion sodium dodecyl sulfate polyacrylamide gel electrophoresis gradient gels and stained with Coomassie Brilliant Blue (Bio-Rad) for normalization of insoluble protein fractions imaged by immunoblot.

### Fluorescence Recovery After Photo-ablation (FRAP)

Day 1 adult worms were used for all FRAP experiments and immobilised on 5% agarose pads for imaging, covered in 2 µl of a solution of 500 µM sodium azide in M9 medium. Coverslips were sealed with molten Vaseline. Imaging was performed in a Nikon A1R confocal microscope with a 100x oil immersion lens (Nikon USA, Melville, NY) which allows for photobleaching of the regions of interest (ROIs). 2 images in 2D were taken pre bleach, and images were taken every 5 seconds for 1 minute post bleach using the 100X oil immersion lens. Two cells or puncta were bleached for every worm. The FRAP measurement was performed with 405 nm Galvano laser at a power of 2. Nikon software reported fluorescent intensity for each ROI at every time point. Loss and return of fluorescent intensity were measured and graphed with GraphPad Prism software, and images were processed in FIJI:imageJ (Brignull *et al*., 2006; Garcia *et al*., 2007; Devkota and Pilon, 2018).

## Acknowledgements

We thank the reviewers for helpful comments and suggestions. We thank Heather Currey, Elaine Loomis, Asia Beale, Lisa Chang, and Hadi Hutchin for outstanding technical assistance. We thank the Developmental Studies Hybridoma Bank (NICHD) for the β-tubulin antibody E7. Some strains were provided by the CGC, which is funded by NIH Office of Research Infrastructure Programs (P40 OD010440), and the National Bioresource Project (Japan). Illustrations Created with Biorender.

## Data and Resource Availability

All relevant data and details of resources can be found within the article and its supplementary information.

## Author Contribution Statement

R.R. performed experiments, analyzed data, edited the manuscript and wrote the first draft manuscript. A.S performed experiments, provided methodology, generated strains, and edited the manuscript. K.P. performed experiments and edited the manuscript. M.H. performed experiments and edited the manuscript. B.C.K. conceived and oversaw the study, analyzed data, and wrote the manuscript.

## Supplemental Figures

**Supplemental Figure 1.**
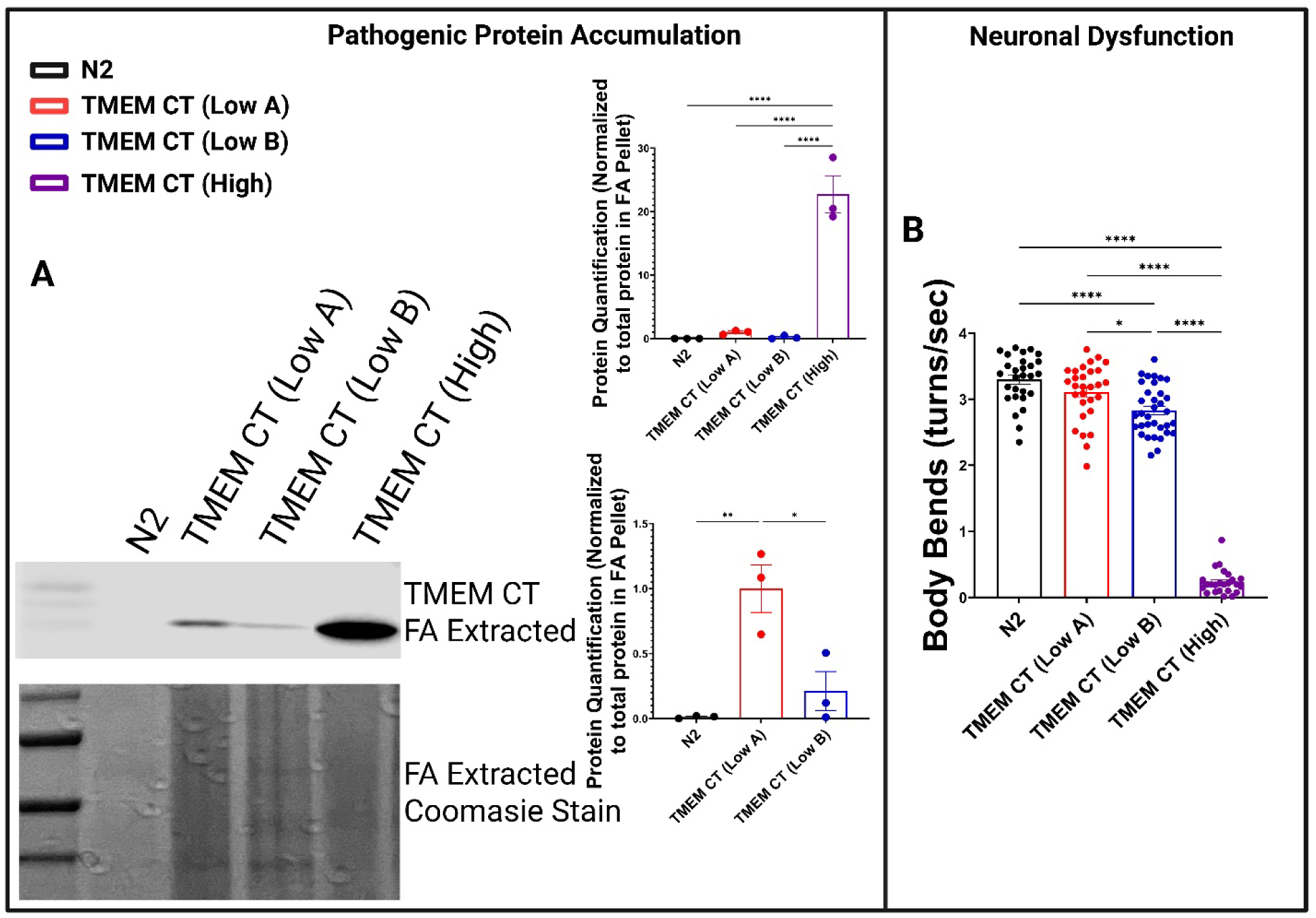
Protein accumulation and neuronal function of low expressing TMEM CT *C. elegans* strains. Comparison of non-transgenic *C elegans* (N2), low expressing transgenic *C elegans* (TMEM CT (Low A) or TMEM CT (Low B)), and high expressing transgenic *C. elegans* (TMEM CT (High)). **A)** Western blot analysis and quantification of *C. elegans* populations against TMEM CT (SYSY TMEM106B – 506 017) for detergent insoluble / FA treated samples. N = 4 for each strain, * = P < 0.05, ** = P < 0.001, **** = P < 0.00001. **B)** Liquid thrashing assay of motor neuron function at day 1 of adulthood, measuring body bends for each worm over a 60 s period. n > 30 for each strain, * = P < 0.05, **** = P < 0.00001.

**Supplemental Figure 2.**
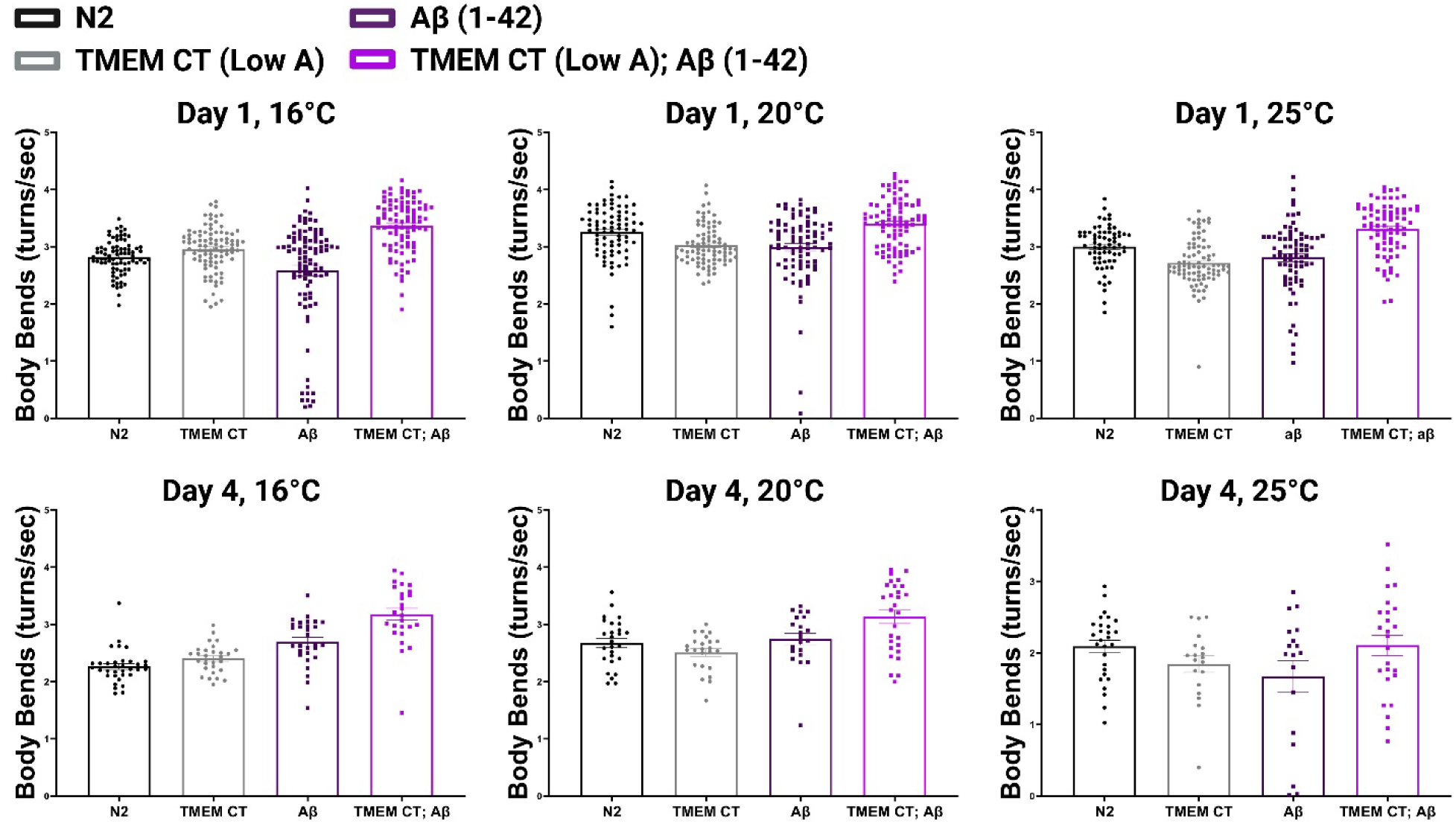
TMEM CT aggregation does not promote neuronal dysfunction synergistically with Aβ_1-42_. Comparison of non-transgenic *C elegans* (N2), single transgenic *C elegans* (TMEM CT (Low A) or Aβ_1-42_), and bigenic *C. elegans* (TMEM CT (Low A); Aβ_1-42_). Liquid thrashing assay of motor neuron function at day 1 of adulthood, measuring body bends for each worm over a 60 s period. n > 60, N = 3 for each strain. Liquid thrashing assay of motor neuron function at day 4 of adulthood, measuring body bends for each worm over a 60 s period. n > 20 for each strain.

**Supplemental Figure 3.**
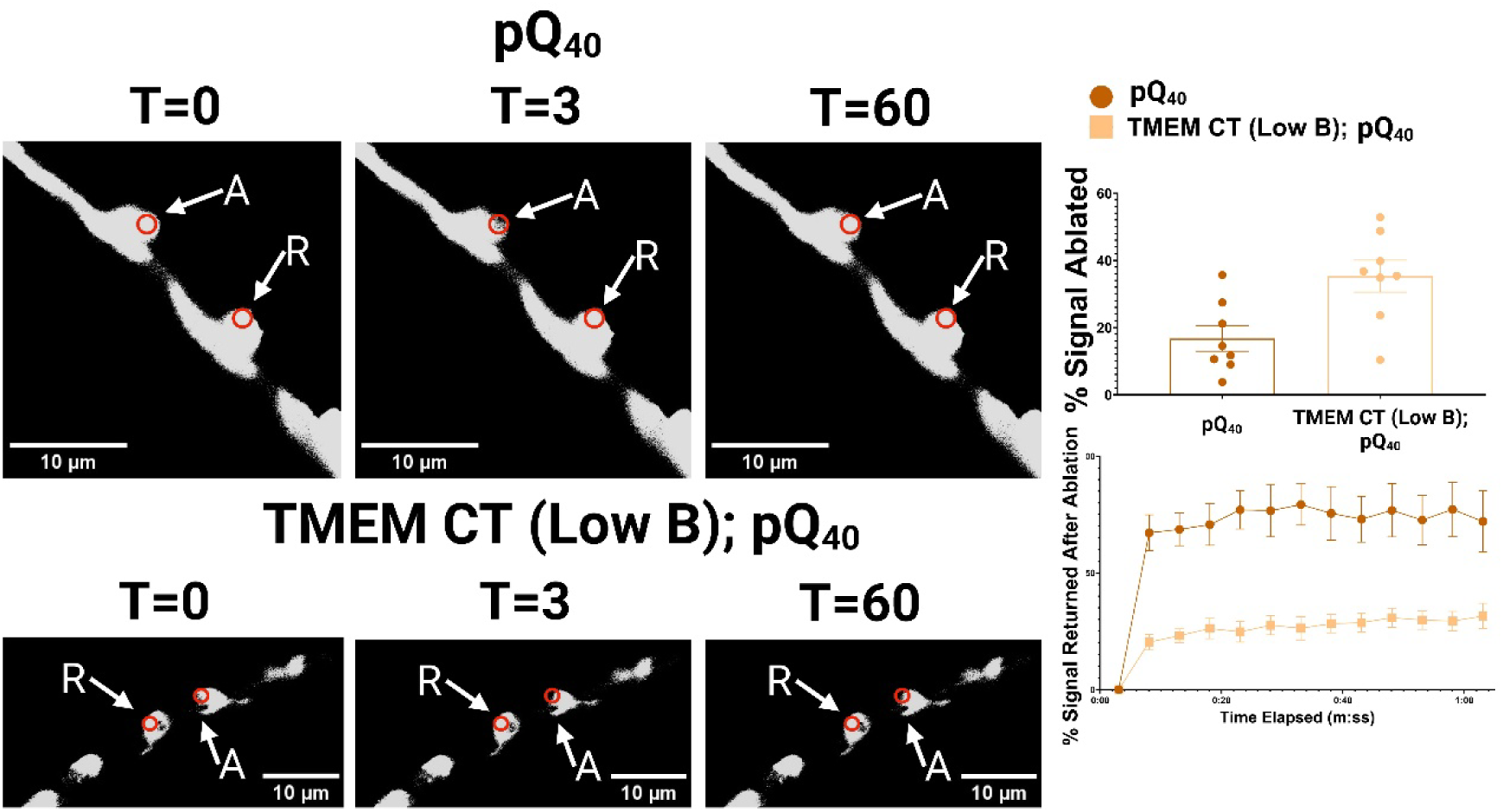
TMEM CT aggregation decreases mobility of polyQ_40_. **(Left)** 100 X magnification of ablated and reference cells or puncta. The left column represents images before ablation, the middle column represents images immediately after ablation, and the right column represents images 1 minute post ablation. Size bar represents 10 µm. Images taken by Nikon Confocal Microscope and processed in FIJI:imageJ. **(Top Right)** Observation of % signal reduction by photobleaching comparing YFP signal of pQ_40_ to TMEM CT (Low B); pQ_40_ (n = 6-8 cells, error bars represent SEM). **(Bottom Right)** FRAP analysis comparing recovery of YFP fluorescence after photobleaching. Red circles indicate ablated or reference areas.

